# Priority effects drive fungal and nematode emergence from insect larvae

**DOI:** 10.1101/2025.11.14.688296

**Authors:** Amaury Payelleville, Magdalena Warren, Chloe Golde, Devyn Sasai, Benny Pan, Patrick Cleary, Arizbel Gomez, Rubina Shrestha, Jean-Claude Ogier, Sophie Gaudriault, Christine Jacobs-Wagner, Tadashi Fukami

## Abstract

Priority effects, in which species arrival history influences community assembly, are increasingly recognized to affect host–parasite systems. However, priority effects across disparate groups of parasitic organisms are poorly understood despite the wide range of taxonomic groups involved. In California oak woodland, we investigated how priority effects between two insect-parasitic fungi (*Metarhizium* and *Beauveria*) influenced emergence of nematodes from insect larvae. Field and laboratory results indicated that both fungi were common, but priority effects prevented them from co-emerging from the same larva. *Metarhizium*- and *Beauveria*-infected insects did not differ in the species composition of emerging nematodes, but larvae without fungal emergence had distinct nematode communities, with *Oscheius* almost always emerging without fungi. Experiments indicated that none of the commonly found nematodes (*Acrobeloides, Mesorhabditis*, *Oscheius*, and *Rhabditis*) were entomopathogenic, but that *Oscheius* could exclude *Beauveria* if arrived early. This time-dependent exclusion was likely caused by an *Ochrobactrum* bacterium that *Oscheius* nematodes carried. Together, these findings suggest that fungi enter insects as primary arrivers, while nematodes come as secondary arrivers to exploit fungus-killed insects, with priority effects influencing both groups. We suggest that this system is a promising natural microcosm for understanding priority effects across disparate groups in host–parasite systems.

## Introduction

It has been a century since Alvar Palmgren articulated the hypothesis that the order and timing of species arrival dictate how species affect one another and, consequently, which species dominate in local communities (Palmgren 1926). Known today as priority effects (Fukami 2015; Drake 1990; Chase and Leibold 2003), this historical contingency in community assembly has been observed in a variety of organisms (Stroud et al. 2024; Grunberg et al. 2023). Among them, coinfecting parasites and pathogens are some of the organisms for which priority effects have been increasingly studied over the past decade (Karvonen et al. 2019).

In studying priority effects, natural microcosms, defined as “small contained habitats that are naturally populated by minute organisms,” have played a central role in advancing basic understanding (Srivastava et al. 2004). A wide range of systems have been used as natural microcosms, and one that has been indicated only recently for their potential consists of entomopathogenic fungi in soil-dwelling insects. As suggested by Costantin et al. (2025), this system can make unique contributions to understanding host–parasite interactions when multiple parasitic species coinfect the same insect.

Costantin et al. (2025) provided evidence for priority effects between two major genera of insect-parasitic fungi, *Metarhizium* and *Beauveria*. When both were experimentally introduced, whichever genus was introduced first emerged more abundantly. However, fungi are not the only organisms that use insect larvae as resources. Soil-dwelling nematodes and their associated bacteria can also establish and emerge from dead insect larvae. Consideration of these other organisms can broaden this system’s scope as a promising natural microcosm. However, few studies have documented how soil-dwelling nematodes or their associated bacteria may interact with *Metarhizium* or *Beauveria* in insect cadavers.

In this paper, we examine the possibility that entomopathogenic fungi *Metarhizium* and *Beauveria* in an insect larvae influence not only each other through priority effects, as previously demonstrated, but that these priority effects influence the community of nematodes that establish and multiply in the insect cadaver. If this possibility is plausible, then that would indicate that priority effects between *Metarhizium* and *Beauveria* could have broader consequences, affecting not just their own performance, but also the community assembly of nematodes in the infected insects. To study this possibility, we conducted a year-long field survey in California oak woodland. We also did a series of laboratory coinfection experiments using the fungi, nematodes, and nematode-associated bacteria that we isolated from the field survey. Based on the findings from this study, we will propose hypotheses on how priority effects among fungi, nematodes, and bacteria influence fungal and nematode emergence from infected insects.

## Materials and methods

### Field survey

#### Study site

We conducted a year-long field survey to assess the presence of active entomopathogenic and opportunistic organisms in the soil beneath coast live oak (*Quercus agrifolia*) trees at Jasper Ridge Biological Preserve (’Ootchamin ‘Ooyakma). In this preserve located in the Santa Cruz Mountains of California in the United States, adults of the acorn moth (*Cydia latiferreana*) and the filbert weevil (*Curculio occidentis*) lay eggs in coast live oak acorns while acorns are still attached to the tree. The larvae hatch and grow within the acorn, and emerge out of the acorn after it falls to the ground in the autumn (Lewis 1992). The emerged larvae then burrow into the soil, which is the period when they can be infected by entomopathogens before they pupate and leave the soil as adults (Bruck and Walton 2007). Given their high abundance (Lewis 1992), these insects likely serve as the main resources for insect-parasitic fungi and nematodes in this preserve.

#### Soil sampling

Soil samples were collected from two different locations within the preserve in February, April, July, and September of 2023 and January and March of 2024. One location (hereafter called location 1; 37.4071868, −122.2344094) was on top of the ridge in *Q. agrifolia* woodland. The other location (hereafter called location 2; 37.4034912, - 122.2347112) was in a small valley that had wetter and more heterogeneous woodland than location 1. At each location, three *Q. agrifolia* trees were haphazardly selected. For each tree, five soil samples, each of 10 cm diameter and a depth of 0-25 cm from the surface, were collected within a two-meter radius from the base of the tree. The five soil samples collected around each tree were then mixed and sieved through a 0.5-cm mesh sieve to remove rocks, roots, and leaves, and then placed in plastic bags. Once transported to the laboratory within two hours of soil collection in the field, the three samples were mixed to make one larval trap corresponding to the respective location.

#### *Ex situ* baiting

Approximately 1 kg of soil from each location was placed inside glass boxes (17 cm wide, 22 cm long, and 6 cm deep) covered with a rubber lid pierced to allow air flow. Since larvae of the *Cydia* moth and the *Curculio* weevil were available for collection only during the few months of the year when they come out of the acorns and burrow into the soil, we used two surrogate species as larval traps: the greater wax moth (*Galleria mellonella*) and the mealworm beetle (*Tenebrio molitor*). These well-studied insects were readily available in large quantities from a vendor (Carolina Biological Supply) throughout the year.

For *Galleria*, 30 fifth-instar larvae were added to each trap, and survival monitored daily for two weeks. When all larvae were dead or when they had been in traps for more than two weeks, all remaining larvae were removed, and a new batch of 30 larvae were added and monitored the same way. The same protocol was followed with *Tenebrio* but using smaller traps (approximately 500g of soil in boxes that were 14 cm wide, 18 cm long, and 4.5 cm deep) and smaller batches of larvae (20 larvae). For both *Galleria* and *Tenebrio*, dead larvae were removed from the trap and added to separate “White traps” (White 1927) containing approximately 20 ml of Ringer’s solution. Each trap was labelled with the larval ID corresponding to the type of larva, the month and location from where the soil was taken, and the date of larval death. The traps were then transferred to an incubator set at 25 °C. After three to four weeks of incubation, larval cadavers in the White traps were checked to determine the presence of fungi, nematodes, and other organisms.

In addition, to test for the effects of soil moisture on fungal and nematode emergence, the moisture level of soil from February 2023 at each location was measured by weighing a soil sample before and after two rounds of dry autoclaving and placement in a dry cabinet for two days. The difference in weight was determined to estimate the amount of water in the soil for the time point of February 2023, the wettest month of the year. For each other month, the same protocol was followed, and the amount of sterile water needed to match the level of moisture observed in February 2023 was added to the soil.

#### Identification of emerging fungi

When fungi were found in a White trap, they were re-isolated on Potatoes Dextrose Agar medium (PDA) plates to obtain a single morphotype. In some cases, several re-isolations were necessary to separate a single morphotype. Two most abundant morphotypes on each plate were identified by DNA extraction with the ExtractNAmp kit (Sigma) and amplification and sequencing of the ITS region with the primers EPF-ITS1-F and EPF-ITS4-R (**Table S1**). The two sequenced isolates were then used for all experiments. All other isolates corresponding to each morphotype were stored but not used in the pathogenicity assay described below, unless stated otherwise.

#### Identification of emerging nematodes

When nematodes were detected in Ringer’s solution of the White traps, they were washed using tap water, sifted by a 20-µm meshed sieve, and then stored at 15 °C in a cell culture flask containing fresh Ringer’s solution for later amplicon sequencing. For each nematode emergence, approximately 1,000 infective juveniles (IJs) were taken from the flasks and frozen at −80 °C in Fastprep tubes when emergence was dense enough. Otherwise, 100 µl containing as many nematodes as possible was taken and frozen.

#### DNA extraction

On the day of the DNA extraction, the tubes were heated to 80 °C and centrifuged at 2000 r.c.f. for 10 minutes. Liquid was then removed, and the nematodes resuspended in 100 µl of the extraction solution from the ExtractNamp kit. Three 2-mm beads were added to each tube. Then, 25 µl of Tissue prep solution (from the above kit) and 20 µl of Proteinase K were added to each tube, and bead beating was performed for 45 seconds after the tubes were incubated at 55 °C for 1 hour. Temperature was then increased to 95 °C for 3 minutes and 100 µl of Neutralizing buffer was added.

#### Amplicon library preparation

Four µl of the DNA extraction solution was used as DNA matrix for the PCR amplification using primers MB_ITS_APJC_F and MB_ITS_APJC_R (**Table S1**) at 0.5 µM final concentration each and using the red ready-Mix PCR. We performed PCR as follows: 95 °C for 3 minutes, followed by 10 cycles of 95 °C for 15 seconds, 52 °C for 30 seconds, 72 °C for 2 minutes, and 30 cycles of 95 °C for 15 seconds, 60 °C for 30 seconds, 72 °C for 2 minutes, ending with 72 °C for 5 minutes. The PCR product was then migrated on an agarose gel to verify amplification. Each PCR product showing clear amplification on the gel was kept. If no amplification was observed, the protocol was repeated.

An amplicon library was prepared for a total of 432 PCR products obtained. Each PCR product was purified using beads (Canvax, AN36, High purity clean-up magnetic beads). Ten µl of beads was added to each well of 96-well plates mixed by pipetting up and down ten times and left on the bench for 5 minutes. Plates were then transferred to a magnetic rack and the liquid removed once all beads were stuck to the side of the well. Two washes using 70% ethanol were then performed, and the plate was left to dry for 5 minutes. Once no ethanol could be detected, 50 µl of 55 °C DNAse-free water was added to each well. Each purified PCR product was then barcoded using the corresponding primer listed (**Table S2**). Five µl of each purified PCR product was added to a new well with 1.6 µl of premixed primers, 10 µl of red ready-Mix PCR, and 3.4 µl of water. The barcoding was performed using the following cycle: 95 °C for 3 minutes, 8 cycles of 95 °C for 30 seconds, 55 °C for 30 seconds, and 72 °C for 30 seconds, ending with 72 °C for 5 minutes.

#### Sequencing

After barcoding, PCR products were migrated on a gel to confirm the intensity of the band and then purified using the same protocol as above, except for the elution volume, which ranged from 20 to 60 µl, depending on the intensity on the gel (higher volume for a less intense band). The samples were then pooled together and sent to the Stanford Genomic Sequencing Service Center, where the concentration and quality of the pooled samples were analyzed with a Fragment Analyzer (Agilent, Santa Clara, CA, USA) and they were then sequenced on a MiSeq (Illumina) using a 2 x 300 cycle sequencing kit with a 15% PhiX spike-in.

Raw BCL files from the MiSeq run were demultiplexed with the Illumina bcl2fastq software, using default settings except for barcode-mismatches, which was set to 0. Index barcodes and Illumina sequencing adaptors were excluded in the resulting fastq files. The forward and reverse primers were removed from the fastq files with cutadapt (Martin 2011). The trimmed sequences were processed with DADA2 (Callahan et al. 2016), with the following modifications: in the filterAndTrim function, truncQ was 2, minLen was 50, and trimRight was 141. The quality of many of the reads deteriorated after about 160 bp, and since our reads were about 300 bp long, we trimmed the right 141 bp to remove the low-quality ends. Since we sequenced the ITS region, the amplicon reads were not long enough to span the entire gene. Therefore, the rest of the downstream analysis was completed with only the forward reads that had better quality than the reverse reads.

#### Sequence data processing

To classify the resulting amplicon sequence variants (ASV), we built a custom BLAST database that included all ITS1, ITS2, and 18S sequences in the Nematoda phylum available in the NCBI nucleotide database. To build the database, we downloaded all the relevant sequences, deduplicated them with seqkit (Shen et al. 2016), and created a database from the deduplicated sequences with the BLAST makeblastdb tool. We then did a blastn search of our ASV sequences in our custom database with a minimum percent identity threshold of 80 for all hits. ASVs that had no hits against our custom database were classified using a blastn of the entire NCBI nucleotide database. The hit with the highest bitscore was retained for each ASV, and taxonomic details were extracted from the hit’s accession lineage. This taxonomy was combined with the DADA2 output count data and sample metadata for downstream analysis with phyloseq. For clearer and more relevant results, ASVs were aggregated at the genus level.

#### Analysis of nematode species composition

Binary presence/absence matrices were constructed with individual insect larvae as rows and identified nematode taxa as columns. Downstream analysis included only identified taxa, excluding non-informative and unidentifiable taxa. Metadata such as treatment group, emerging fungal morphology, and concentration were included in the analysis.

We calculated a Jaccard dissimilarity matrix of the presence/absence ASV matrix using the vegdist function from the vegan package (Oksanen, 2012). Principal Coordinates Analysis (PCoA) was then performed on the dissimilarity matrix with the pcoa function from the ape package (Paradis and Schliep 2019). The first two axes were extracted for visualization of sample clustering based on emerging nematode species composition.

To evaluate whether groupings in species composition were statistically significant, a permutational multivariate analysis of variance (PERMANOVA) was conducted using the adonis2 function (999 permutations). This analysis tested for differences between treatment groups or morphology types based on the Jaccard dissimilarity matrix. To assess whether significant PERMANOVA results were driven by variation in group dispersion, a multivariate homogeneity of dispersion test (PERMDISP) was conducted using betadisper followed by anova.

To identify which environmental or treatment-related variables were significantly associated with variation in nematode species composition, environmental vectors were fitted to the PCoA ordination using the envfit function. The analysis tested the correlation of factors (e.g., fungal morphology, treatment, concentration) with the PCoA axes. Significance was evaluated using 999 permutations. Only variables with p values < 0.05 were interpreted as meaningfully aligned with community variation.

After running the PCoA, the coordinates of the first two axes for each sample were extracted and grouped depending on fungi presence/absence. The mean coordinates of each axis (centroids) for each group (*Beauveria*, *Metarhizium*, and no fungus) were then calculated and represented in the PCoA plot.

The optimal number of clusters of the nematode ASVs, in this case seven, was calculated by computing the within-groups sum of squares for the clusters produced by the k-means method. Hierarchical clustering analysis of the Jaccard distance matrix was conducted with the hclust function, and the result was cut into seven groups using the cutree function. The clusters were visualized with a dendrogram using the dendextend and ggdendro packages (Galili 2015; de Vries and Ripley 2024).

#### Fungal pathogenicity on *Galleria* larvae

*Beauveria* and *Metarhizium* fungi were grown on PDA for one week or until spores covered the plate. Ten ml of Ringer’s solution was then added on top of the plate, and the plate was gently scratched with a 1-µl loop to resuspend spores. The Ringer’s solution was then transferred to a 50-ml Falcon tube, and this process repeated once. Concentration of spores was then measured using a hematocytometer. Volume was adjusted to reach an equivalent concentration for different fungi. After thorough vortexing, larvae were immersed in the spore solution and individually put in wells in 12-well plates, and larval survival was monitored at 25 °C over time. Larvae used as the control were immersed in Ringer’s solution without spores. Each experiment was conducted with 20 larvae per treatment and repeated two to four times, depending on the availability of spores and larvae.

Larval survival following exposure to *Beauveria* or *Metarhizium* spores was analyzed using a binary status variable, i.e., death or survival, to compare three treatment groups: control, 10 spores/ml, and 10 spores/ml. Kaplan–Meier survival curves were constructed using the survfit() function from the survival R package. Survival probability over time was estimated separately for each concentration group within each fungal treatment (*Beauveria* and *Metarhizium*). Differences between survival curves were evaluated using the log-rank test, as implemented in ggsurvplot() from the survminer package.

To quantify the effect of fungal spore concentration on larval mortality, we fitted Cox proportional hazards models using the coxph() function. The model included spore concentration as the predictor, with 10 spores per ml as the reference level. Hazard ratios and 95% confidence intervals were extracted from the model summaries. A hazard ratio greater than one indicated an increased risk of death relative to the reference group, while a value less than one indicated a protective effect. Statistical significance of the model terms was assessed using Wald tests, and overall model fit was evaluated using three global tests: likelihood ratio test, Wald test, and score (log-rank) test.

#### Nematode pathogenicity on *Galleria* larvae

Concentration of nematodes in Ringer’s solution was measured by counting under a dissecting microscope. A solution containing estimated 1,000 nematodes/100 µl was prepared to assess pathogenicity. For inoculation, we used 2-ml Eppendorf tubes containing a piece of filter paper folded to cover the sides of the tubes, with the lid pierced to allow air flow. One *Galleria* larva was added to each of these Eppendorf tubes and then 100 µl of the nematode solution was added to the filter paper inside the tube. Survival of the larvae was then monitored over time at 25 °C, repeated for 20 independent larvae. For some nematodes, no biological replicate for each isolate was possible due to a low number of nematodes available. Results for replicates for other isolates can be found in **Table S3** and correspond to a batch of 10 *Galleria* larvae each.

#### Interactions between fungi on *Galleria* larvae

To study interactions between fungi, insect larvae were dipped into Ringer’s solution with varying amounts of *Beauveria* and *Metarhizium* spores (10^5^, 10^4^, or no spores). Larval survival was monitored over time. Fisher’s exact tests were conducted between the following treatment pairs of *Beauveria* / *Metarhizium*: 10^5^/10^5^ vs. 10^5^/10^4^, 10^5^/10^5^ vs. 10^4^/10^5^, and 10^5^/10^4^ vs. 10^4^/10^5^.

#### Interactions between fungi and nematodes on *Galleria* larvae

To study interactions between fungi and nematodes, insect larvae were first dipped into Ringer’s solution with fungal spores (either 10^5^ or 10^4^ spores) and then transferred to Eppendorf tubes containing a piece of filter paper with 1,000 nematodes. Larval survival was monitored over time. A Chi-squared test of independence was conducted on a contingency table of treatment conditions versus emerging organism identity to evaluate whether the distribution of emerging organisms differed significantly across co-infestation treatments.

#### Isolation of bacteria from crushed nematodes on agar plates

To study how nematode-associated bacteria interact with *Beauveria*, we first isolated bacteria from some of the nematodes that emerged in the field survey described above (**Table S4)**. To this end, 1 ml of Ringer’s solution containing emerged nematodes was washed using a 20-µm mesh sieve and added to an Eppendorf tube. After centrifugation at 8,000 r.c.f. for 5 minutes, the supernatant was removed, and the nematode pellet was crushed using a sterile pellet pestle. Bacteria were then streaked on NBTA plates (with a ten time lower concentration of TTC than usual recipe to allow more bacteria to grow) using a 1-µl inoculation loop. After isolation of each colony morphology on the NBTA plates, we amplified and sequenced the *rpoB* gene (Ogier et al. 2020a) of the bacteria to identify bacterium’s genus and species when possible.

#### Interactions between fungi and bacteria on plates

Using the isolated bacteria, we did *in vitro* experiments to study their interactions with *Beauveria*. Fungal spores were spread vertically using a 1-µl inoculation loop on a PDA/LB plate (mix of 18.5 g of PDA with 13 g of LB agar in 1000 ml of water). After growing each bacterial isolate independently in 5 ml LB broth overnight, each isolate was streaked horizontally on the PDA/LB plate where the fungal spores had been streaked. Three different bacterial isolates were added for each plate. Each species combination was conducted at least nine times. Plates were incubated for a week at 25 °C. Plates were monitored over time and growth of the fungi quantified by the development of mycelium on the plate and similarly growth of bacteria by presence of colonies or viscous streak. A time-delay experiment was conducted similarly except that we streaked *Beauveria* 24 hours before streaking the bacteria on the plate. A pairwise Fisher’s exact test was used to compare the different outcome of each pairwise competition.

#### *In vivo* competition in *Galleria* larvae

We also did *in vivo* experiments, where *Beauveria* spores were applied to *Galleria* larvae using the same protocol as above. For bacterial injection, bacteria were grown overnight in LB broth. Fifty µl of the overnight culture was then transferred to 5 ml of fresh LB broth and grown for 4 hours at 25 °C until it reached an optical density (600 nm) of 0.1 to 0.3. After washing the cells by two rounds of centrifuge (8,000 r.c.f. for 5 minutes), dilutions were made and plated to estimate the number of viable cells injected into each larva. Injection into larvae was done using a microinjector with foot pedal (Chemyx, model F100X). Each larva was injected with 10, 100, or 1,000 bacterial cells. We did fungal spore application 24 hours before bacterial injection (F24/B) and in some cases 48 hours before (F48/B). Each larva was monitored for a week, and cadavers were kept for one more week to monitor fungal emergence. When white mycelia were observed on the cadaver, the emerging organism was considered fungi. When cadavers had no noticeable fungi, the cadaver was cut open and some of the tissue inside was streaked on NBTA plates. Bacteria resulting from this streaking had the same morphology as the injected bacteria. Thus, we assumed that all larva without fungi had bacteria as the emerging organism. As no major difference in the outcome was noticed depending on bacterial concentration, the data were pooled for statistical analysis.

## Results

### Fungal and nematode taxa showed specific associations with one another in insect larvae

Fungi and nematodes were the two major groups of organisms emerging from *Galleria* larvae in February 2023 in soil sample from location 1 (**Figure 1A**). Sixty-one cadavers (44%) yielded nematodes alone, 38 (27%) yielded fungi alone, 30 (21%) had both, and 10 (7%) contained other organisms we could not identify (**Figure 1A**). Nematode emergence declined sharply in April (5 cadavers, 6%), dropped to just one cadaver in July, and disappeared in September. Fungi followed a similar change, though less pronounced. Larval survival increased over this period, with 14% in April, 25% in July, and 35% in September (**Figure 1A**). In January and March 2024, nematodes and fungi reappeared, with nematodes emerging from 37 cadavers (62%) in both months, fungi from 1 (2%) and 3 cadavers (5%) respectively, and both nematodes and fungi from 7 (12%) and 11 (18%) (**Figure 1A**). When we moistened the soil, we detected nematodes in the previously negative samples from July and September (**Figure 1B**), indicating that nematodes were present but probably unable to reach and therefore emerge from larvae due to insufficient moisture. Fungal detection also increased following soil rehydration.

**Figure 1:**
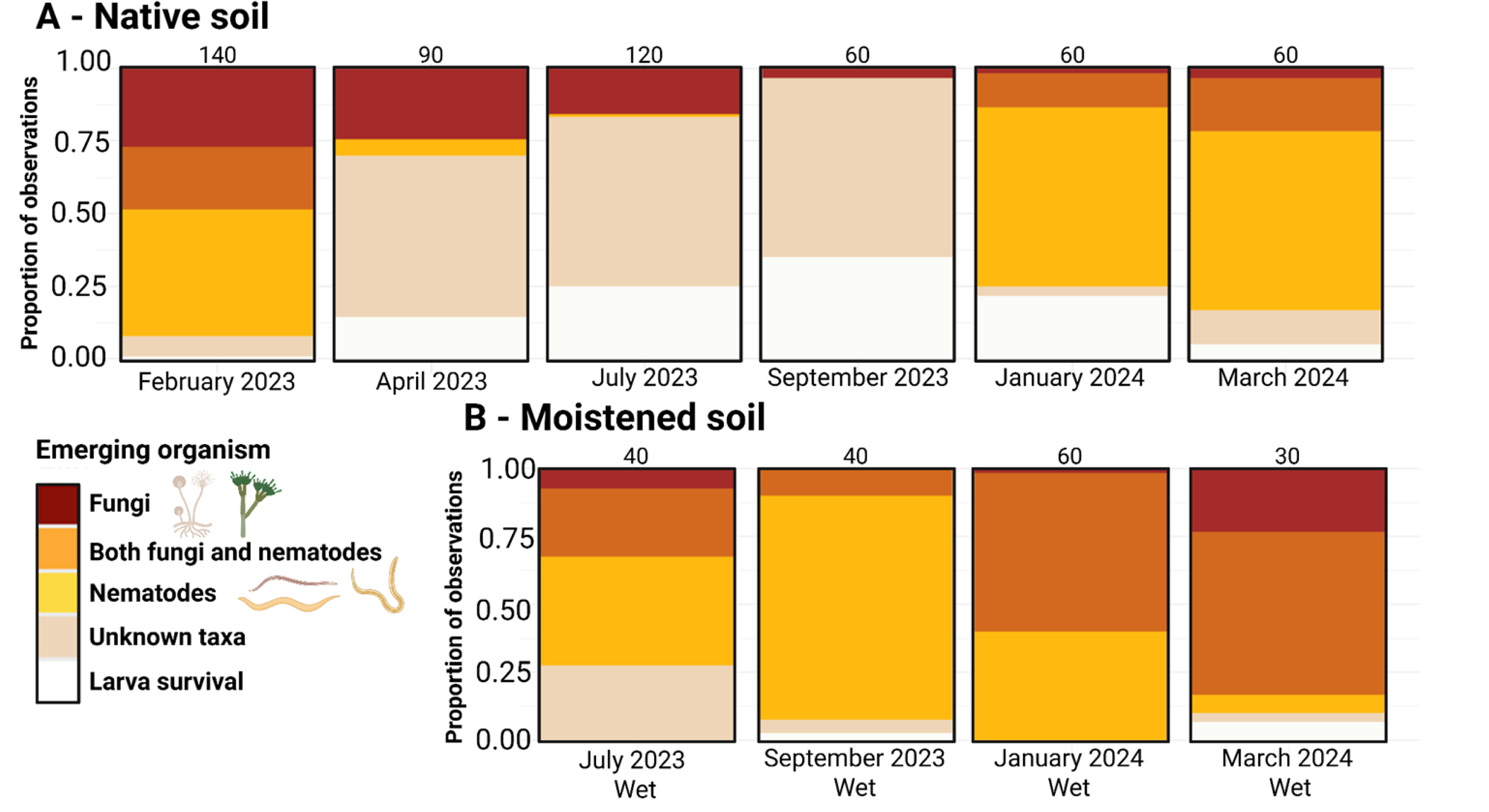
Organisms emerging from *Galleria* moth larvae in the field survey at Jasper Ridge Biological Preserve (’Ootchamin ‘Ooyakma). Fungi and nematodes emerged from *Galleria* larvae placed in the field-collected soil throughout the survey period from February 2023 to March 2024, although the proportion of fungal, nematode, and co-emergence varied from month to month. The field collected soil was used without (A) or with (B) water added to the soil to make the soil as moist as it was in the field during the wettest month (February 2023). Each bar represents the proportion of emerging organisms for that month. The number of *Galleria* larvae tested in each month is shown above the bar corresponding to the month.

In fungi, two distinct morphotypes were common. We sequenced the Internal Transcribed Spacer (ITS) region of the genome of one representative from each, confirming their identity as *Beauveria* (white morphotype) and *Metarhizium* (green morphotype). *Beauveria* was more frequently observed (89 cadavers, 66%) than *Metarhizium* (45 cadavers, 34%). Despite both being common, *Beauveria* and *Metarhizium* rarely co-emerged. Only two of the 132 cadavers with fungal emergence had both (**Figure S1**).

Four nematode genera were commonly detected, emerging either alone or in various combinations (**Figure 2**). Of these genera, *Oscheius*, *Mesorhabditis*, and *Rhabditis* are regarded as entomo-opportunists (Manjula et al. 2020; Zhang et al. 2020; Zhou et al. 2017; Taboga 1981), while *Acrobeloides* is bacterivorous (Postma-Blaauw et al. 2005). In the analysis of nematode species composition, seven distinct clusters emerged (**Figure 2**). *Oscheius*, *Acrobeloides*, and *Mesorhabditis* were the primary genera characterizing nematode community structure.

**Figure 2:**
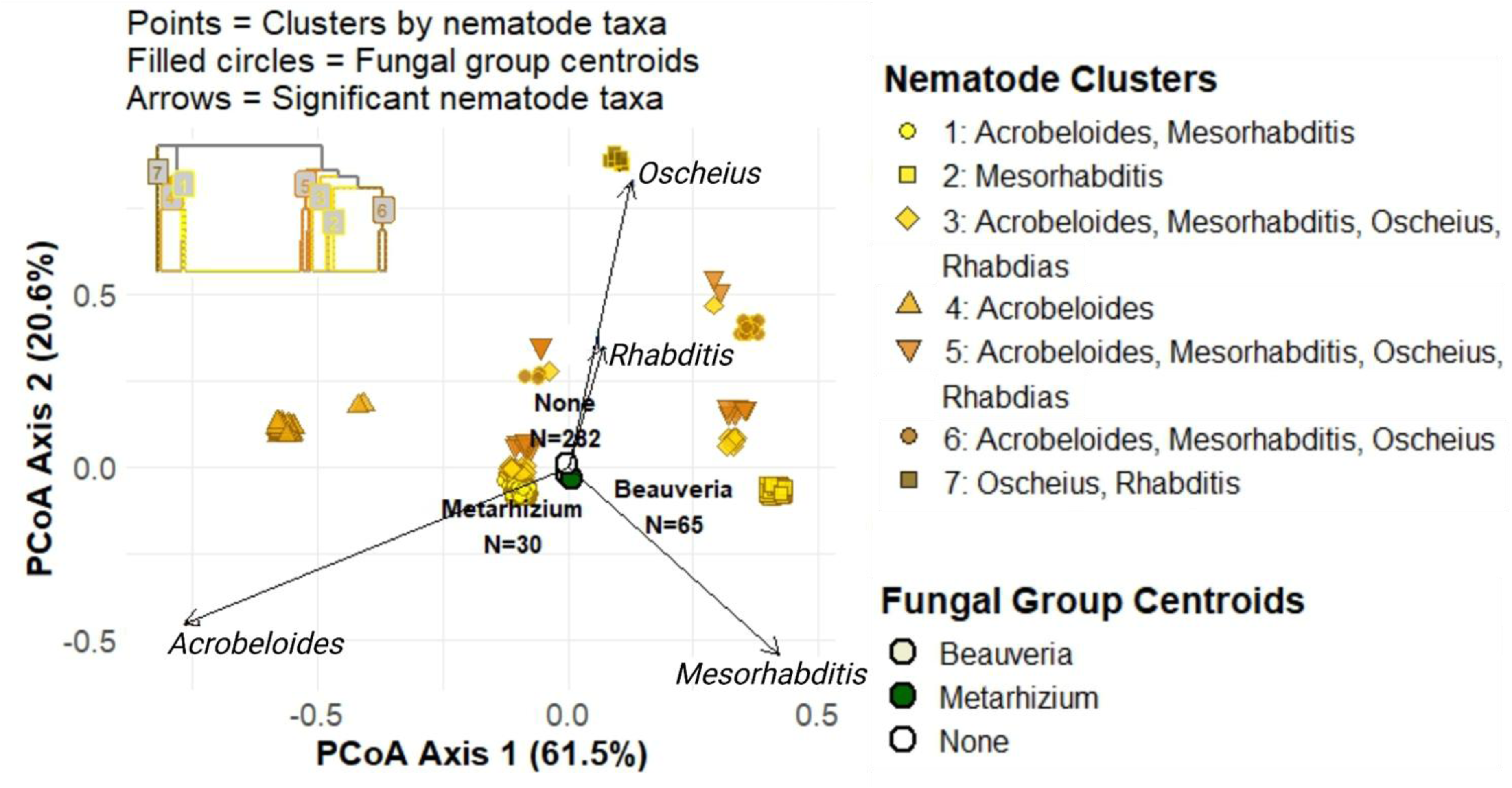
Principal Coordinates Analysis (PCoA) summarizing variation among the *Galleria* moth larvae tested in the field survey in the species composition of nematodes that emerged from the larvae. The PCoA plot is based on the nematode amplicon sequencing data from the field survey conducted at Jasper Ridge Biological Preserve (’Ootchamin ‘Ooyakma) from February 2023 to March 2024, using the same samples shown in Figure 1. Each data point in the PcoA plot represents a larva tested, showing the species composition of nematodes that emerged from the larva. Each point is color-coded according to the cluster group it belongs to in the cluster analysis used to find distinct groups of nematode species composition. Dendrogram summarizing the cluster analysis is shown in the inset. Arrows indicate the contribution of major nematode taxa to each PcoA axis. In addition, the PcoA plot also show the centroids of the larval samples from which *Metarhizium* fungus emerged, those from which *Beauveria* fungus emerged, and those from which no fungus emerged (none), with the sample size for each group shown.

When we added three fungal emergence centroids (*Beauveria*, *Metarhizium*, and no fungus) onto the PCoA summarizing nematode species composition, the centroid positions for *Beauveria*- and *Metarhizium*-emerging larva were close to each other with no statistically significant separation (**Figure 2**). Larvae with no fungal emergence clustered slightly but not significantly separately from *Beauveria*- and *Metarhizium*-emerging larvae. Using the PcoA axis, the four main nematodes genera were clustered highly significantly (p-value < 0.001) separately. Interestingly, *Acrobeloides* was closer to *Beauveria* centroid while *Mesorhabditis* was closer to *Metarhizium*. The most interesting correlation was between *Oscheius* nematodes and the “no fungi emerging” centroid (**Table 1**).

The two sampling locations largely mirrored each other (**Figure S2**). One difference was that fungivorous mites, feeding primarily on *Beauveria*, were present only at the second location and included in the unknown group. Furthermore, the other insect tested, *Tenebrio*, showed higher survival during the dry summer months (**Figure S3**), but we detected similar fungal and nematode communities in *Tenebrio* and *Galleria* during the wet winter months (**Figure 1B and S4**).

### Fungi were entomopathogenic, and exerted priority effects on each other

Both *Beauveria* and *Metarhizium* were pathogenic to *Galleria* larvae (**Figure 3A**). When larvae were exposed to a solution containing 10 spores/ml, the median lethal time (LT_₅₀_) was approximately 70 hours post-infestation for both fungal species. Reducing spore concentration to 10 spores/ml increased LT_₅₀_ to around 160 hours. We isolated and preserved approximately 200 *Beauveria* and 100 *Metarhizium* isolates from this survey. Although only a subset of these strains were tested in the assays reported here, all tested isolates were pathogenic to *Galleria*, with little variation in LT_₅₀_ (data not shown).

**Figure 3:**
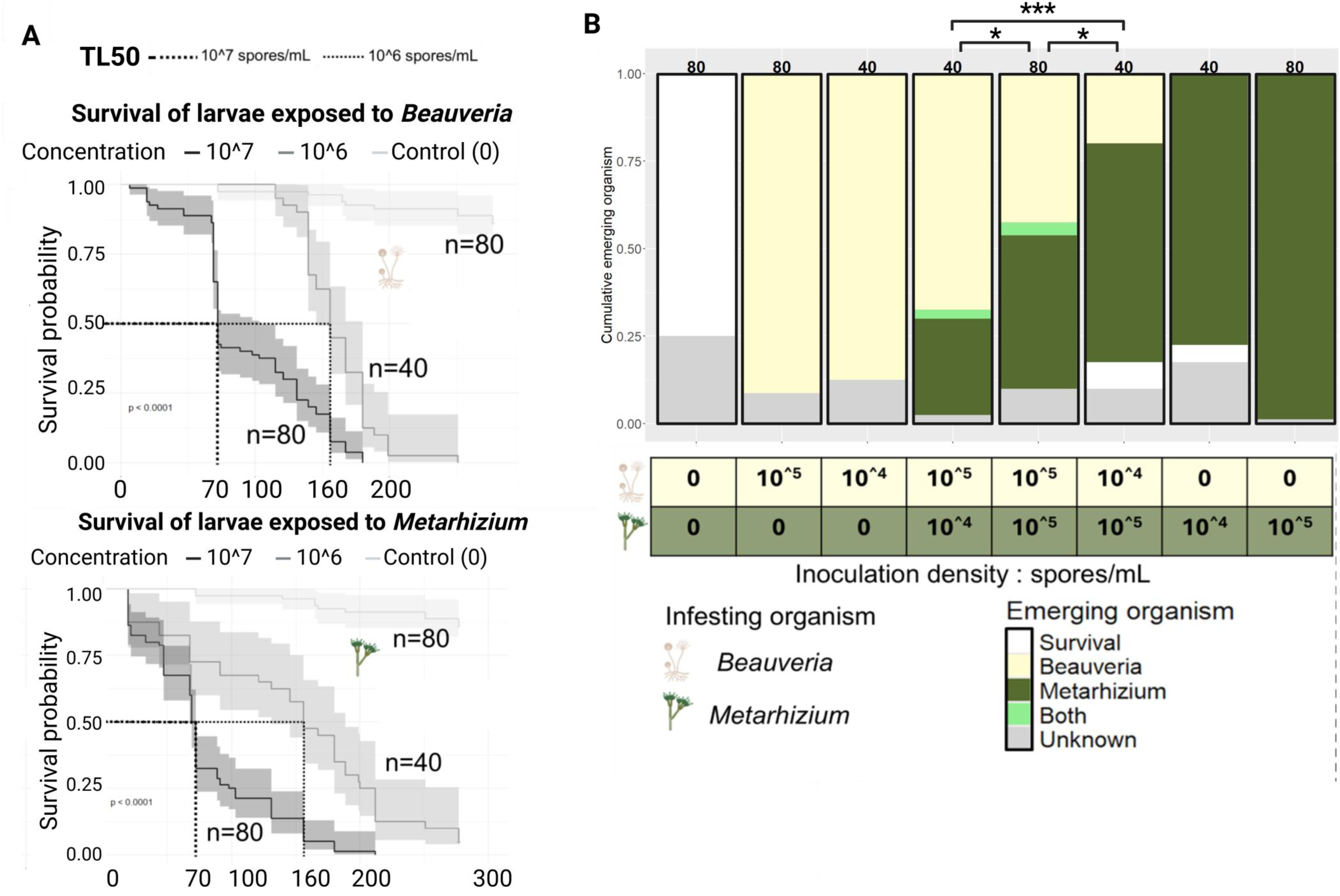
Survival curves of *Galleria* moth larvae that were exposed to a specific species of fungi and priority effect between fungi. (A) the plots show the survival of *Galleria* larvae after exposure to two fungal species (either *Beauveria* or *Metarhizium*) at two spore concentrations (10⁷ and 10⁶ spores/ml), compared with a control exposed to Ringer’s solution. The median lethal time (LT_50_) for each curve is shown by dash line and displayed on x-axis and Kaplan–Meier survival curves are used. (B) *Galleria* larvae were infected by either or both of *Beauveria* (beige) and *Metarhizium* (green) fungi, the plot shows the proportion of larvae from which either species, both species, or neither emerged, at varying spore concentrations used for infection. The concentration of spore/ml used is shown in the table below the graph. Numbers above each bar represent the number of larvae used for each infection treatment. Asterisks indicate results of the statistical comparisons performed using pairwise Fisher’s exact tests (A) where * denotes p <0.05 and *** denotes p < 0.001.

As expected, inoculation with either *Beauveria* or *Metarhizium* resulted in the emergence of the same fungus from the cadaver. In coinfection, only a single fungus emerged in most cases, with only 3% (4 out of 160) of cadavers showing co-emergence (**Figure 3B**). The fungus introduced at a higher concentration was significantly more likely to dominate and emerge, indicating priority effects (Chi-square test, *p* < 0.001; Fisher Exact test for priority effect condition, p-value = 5.385e-05).

### Nematodes were not entomopathogenic, but one genus could outcompete Beauveria *fungi*

None of the nematode isolates caused significant larval mortality (**Figure 4A**). All larvae survived, except in the case of *Oscheius* sp., which resulted in two deaths out of 20 trials. This overall lack of pathogenicity was unexpected, given that nematodes were among the most frequently recovered organisms in our field survey. This finding indicates that the nematodes were not entomopathogenic, despite their frequent emergence from the larvae.

**Figure 4:**
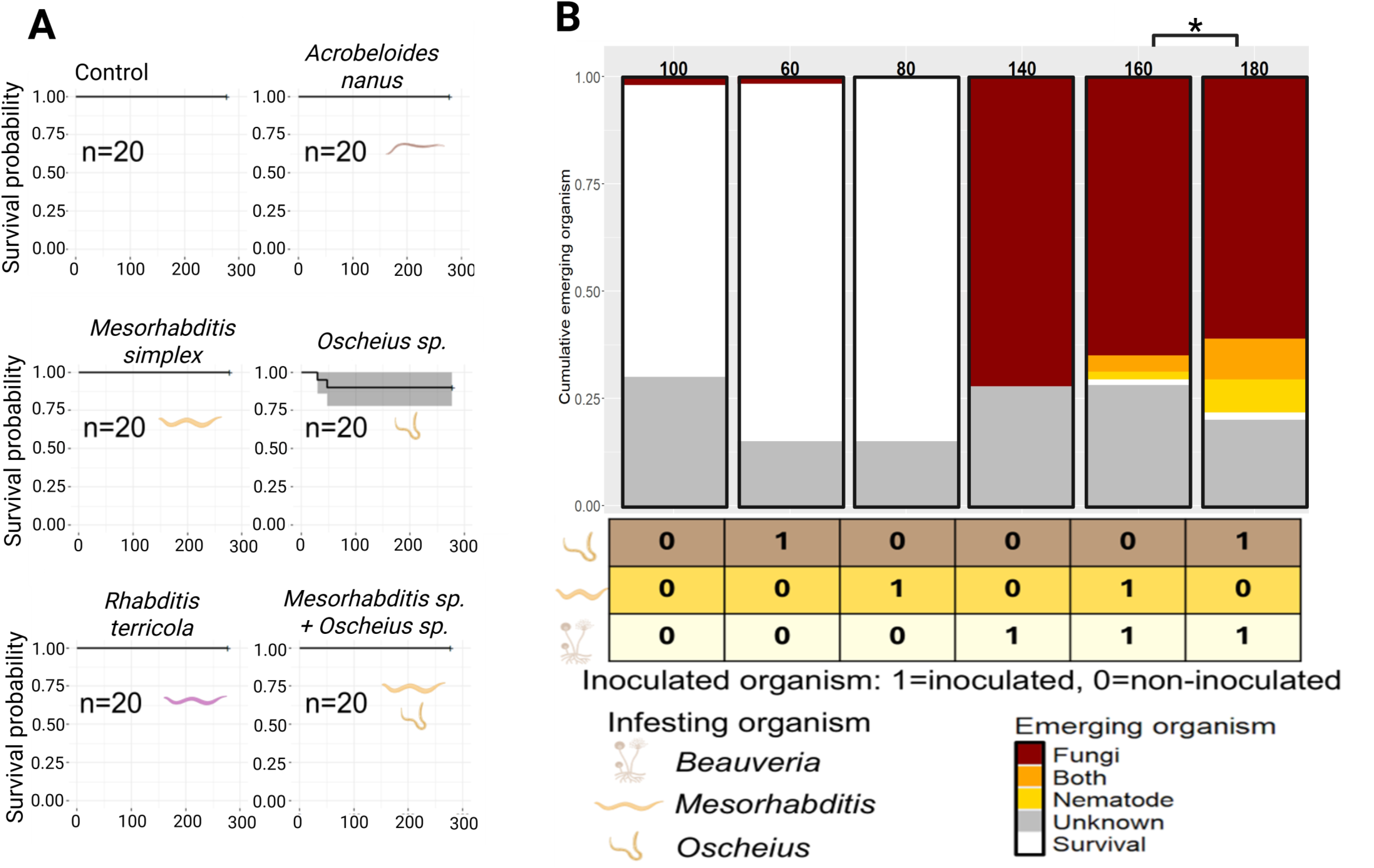
Experimental coinfection of *Galleria* moth larvae by fungi and nematodes. (A) the plots show the survival of *Galleria* larvae exposed to different nematode species at an estimated concentration of 1,000 nematodes per 100 µl of solution, Kaplan–Meier survival curves are used. (B) *Galleria* larvae were infected by *Beauveria* and one of the two species of nematodes, either *Mesorhabditis* or *Oscheius*, the plot shows the proportion of larvae from which the fungus, the nematode, or both emerged when they were infected with 10^^5^ spores/ml for fungi and 1,000 nematodes. Statistical significance was assessed with a chi-squared test of independence. Asterisks indicate results of the statistical comparisons performed using a Chi-square test of independence (B), where * denotes p <0.05.

In coinfection with *Beauveria*, *Mesorhabditis* nematodes emerged from the cadaver in only 6% of cases (10 out of 160). In contrast, coinfecting larvae with *Beauveria* and *Oscheius* resulted in higher nematode emergence rate, with 20% of cadavers (36 out of 180) showing emergence of either nematodes alone or both nematodes and fungi (**Figure 4B**).

### Exclusion of Beauveria fungi by Oscheius nematodes was likely caused by associated bacteria

Sequenced bacteria from nematodes were selected for further study (bold in **Table S4**) based on their detection with only one nematode genus (specialist) or multiple different nematodes genus (generalists). *Ochrobactrum sp.* was the only bacterial species found co-occurring only with *Oscheius* nematodes in our experiments. However, other sequenced bacteria like *Serratia* or *Pseudomonas protegens* were potential specialists based on literature (Zhang et al. 2019; Ogier et al. 2020a; Ruiu et al. 2022) and were therefore selected for the experiments. We used *Achromobacter spanius* (present with all nematodes) and *Ochrobactrum quorumnoscens* (different *rpoB* sequence from the *Ochrobactrum* sp. above and present with both *Oscheius* and *Mesorhabditis* nematodes) as generalists. We added *Enterococcus casseliflavus* isolated from the non-infested *Galleria* larvae, as negative control.

*In vitro* coinfection of selected bacteria and fungi showed two groups of bacteria with respect to their interactions with *Beauveria*. One group, including *Enterococcus casseliflavus*, *Ochrobactrum quorumnoscens*, and *Achromobacter spanius*, rarely outcompeted *Beauveria* when introduced simultaneously with fungi, and never when *Beauveria* had a 24-hour advantage. In contrast, the other group, including *Pseudomonas protegens*, *Serratia* sp., and *Ochrobactrum* sp., almost always outcompeted *Beauveria* even when the fungus had a 24-hour advantage (**Figure 5A**).

**Figure 5:**
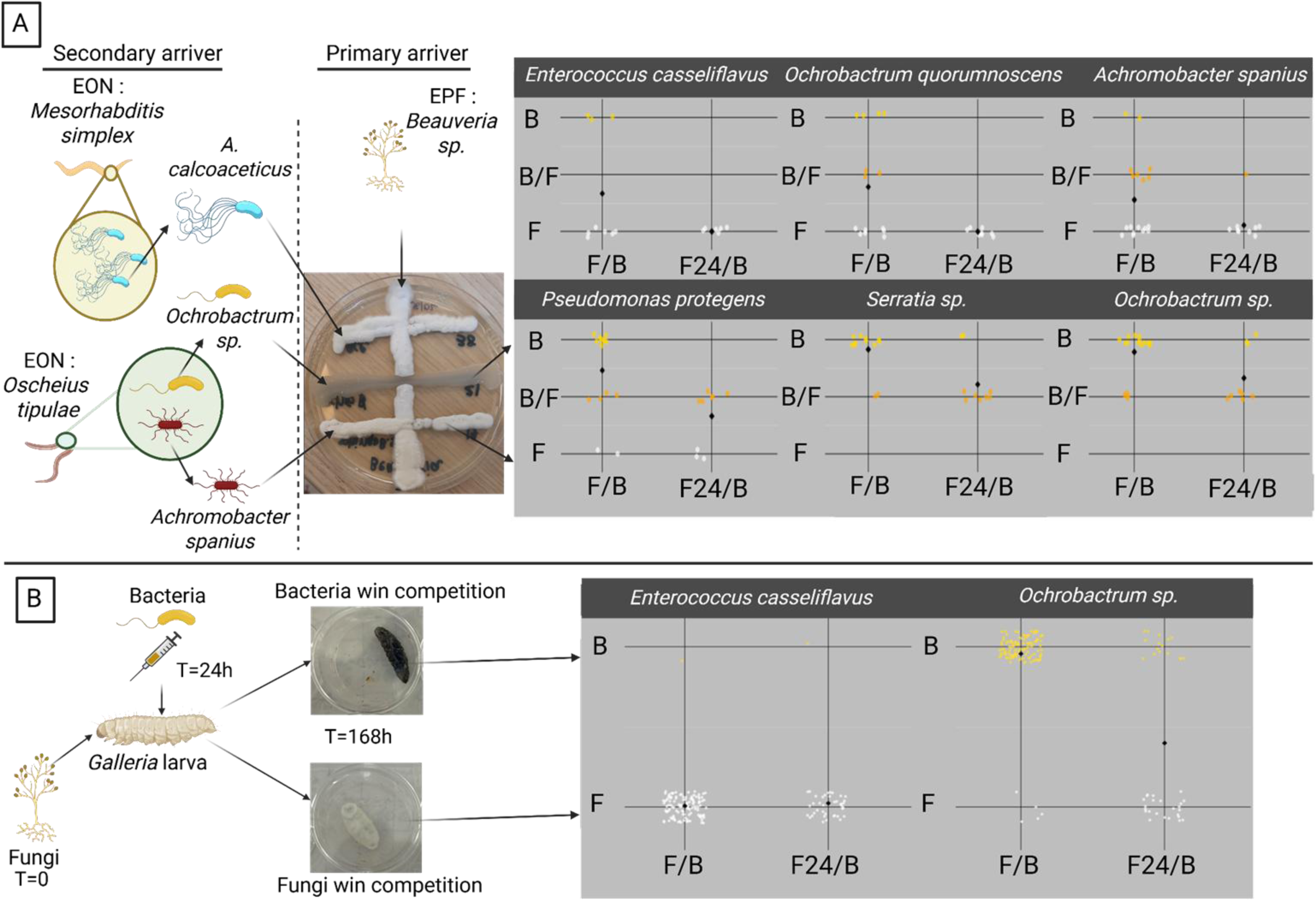
Experimental tests of interactions between *Beauveria* fungus and different bacterial strains isolated from *Galleria* moth larvae and the nematodes that emerged from *Galleria* larvae in the field survey. In *in vitro* experiments (A), bacteria isolated from nematodes or *Galleria* larvae were streaked on Petri dishes with *Beauveria* spores. After growth, if mycelia developed on the whole line, it was counted as the fungus winning against the bacterium (F). If only the bacterium was present, it was counted as the bacterium winning the fungus (B). If both were present, it was counted as a tie (B/F). Results are presented in plots, which also include those of the delayed experiments where the bacteria were introduced 24 hours after streaking of fungal spores (F24/B). In *in vivo* experiments (B), *Galleria* larvae were put in contact with *Beauveria* spores, and either *Enterococcus casseliflavus* or *Ochrobactrum sp*. were injected to the larvae 24 or 48 hours after fungal spore introduction. After larval death, emerging organism was noted (B or F). In both panels A and B, black diamonds on the graphs indicate the mean of the values for each experimental treatment.

These *in vitro* results were consistent with *in vivo* results with *Galleria* larva. When *Enterococcus casseliflavus* and *Beauveria* were put in competition in *Galleria* larvae, only fungi emerged from the cadaver. In contrast, *Ochrobactrum* sp. emerged in most cases without *Beauveria* emergence. Delaying *Ochrobactrum* sp. introduction led to more fungal emergence, although more than half the cadavers were still dominated by *Ochrobactrum* sp. (**Figure 5B**).

## Discussion

Taken together, our results indicate that no clear alternative communities of nematodes exist in a way that corresponds to which of the mutually exclusive fungi, *Metarhizium* or *Beauveria* (**Figure 3B**), emerges from insect larvae. Nematode species composition was indistinguishable between *Metarhizium*- and *Beauveria*-emerging larvae (**Figure 2**). Sometimes only the fungus emerged with no nematodes. Other times the fungus emerged with multiple nematode species (**Figure 2**), but the species identity of these nematodes varied from one larva to another, with no clear distinction between *Metarhizium* and *Beauveria*-emerging larvae. However, the most curious case observed in this study was when nematodes emerged without any fungus (**Figures 1 and 2**). In these instances, the nematodes must have replaced whichever fungus initially killed the insect since no nematode species studied here could reliably establish in a larva by itself (**Figure 4A**). Exploring how fungi interact with not just nematodes, but also bacteria (**Figure 5**), has led us to discover that it was probably inhibition by some bacterial symbionts of the nematodes, especially *Ochrobactrum* sp. in *Oscheius*, that enabled the nematodes to replace the early-arriving fungus.

Most studies investigating entomopathogenic communities have focused on *in vitro* assays using isolated and well-known pathogens (fungi, nematodes, or bacteria) against insect larvae (Lacey et al. 2015; Eleftherianos et al. 2010). Some studies have explored which organisms are present in the soil (Emelianoff et al. 2008; Noujeim et al. 2011; Araújo and Hughes 2016; Topuz et al. 2016; Depuydt et al. 2024; Zhang et al., 2024), but combined use of field survey, experimental validation and investigation of interactions explaining the observations remains rare. In our work, the field survey revealed two main groups consistently emerging from the larvae in this study system in California: fungi, primarily *Beauveria* and *Metarhizium*, and nematodes, mainly *Mesorhabditis*, *Oscheius*, and *Acrobeloides*, (**Figure 1A and 2**). The two fungi rarely co-emerged (**Figure S1**), whereas several nematode species were often observed co-emerging with a fungus **(Figure 2**). Experimental validation showed that fungi were pathogenic to larvae, while nematodes alone caused little mortality (**Figure 3A-4A**). The discrepancy prompted us to explore whether coinfection might explain the high prevalence of nematode emergence. First, we co-infected larvae with both fungal species and confirmed that typically only one fungus would emerge, indicating mutual exclusion through priority effects (**Figure 3B**), consistent with Constantin et al. (2025). As *Oscheius* nematodes were almost never co-emerging with fungi (**Figure 2**) and *Beauveria* and *Mesorhabditis* were the two prevalent organisms of each group in our ecosystem (**Figure S1**), we performed coinfection with one fungus (*Beauveria*) and a nematode (*Mesorhabditis* or *Oscheius*), observing co-emergence in some cases, replicating field observations, but also cases where only the nematode emerged, suggesting it had outcompeted the fungus (**Figure 4B**).

Some nematodes have been proposed to opportunistically use cadavers killed by other pathogens (Blanco-Pérez et al. 2017; Duncan et al. 2003; Campos-Herrera et al. 2015; Blanco-Pérez et al. 2019). To our knowledge, however, this study is the first to demonstrate such behavior between nematodes and entomopathogenic fungi isolated from the same environment. Moreover, a likely explanation for this behavior is given by our finding that some bacteria associated with the nematodes could inhibit fungal growth (**Figure 5**). Some of the bacteria used in our experiment were generalists (*O. quorumnoscens, A. spanius*) in the sense that they were isolated from several species of nematodes and *Galleria* larvae (*E. casseliflavus*). Some other bacteria were specialists in the sense that they were found only when one specific nematode was present (*Ochrobactrum* sp.) or reported in the literature as specifically associated with nematodes (*Serratia* sp. and *P. protegens*) (Ogier et al. 2020b; Torres-Barragan et al. 2011). Comparing how the generalists and the specialists interacted with *Beauveria*, we found no significant difference between bacteria within each group (generalists or specialists), but significant difference was observed between the two groups (**Figure S5**), with the specialists, but not the generalists, having the ability to inhibit fungi. This pattern suggests that only the specialist bacteria associated with specific nematode species may facilitate nematodes in establishing in insects killed by fungi.

Taken together, these findings lead us to classify the organisms emerging from insect larvae into two groups: fungi as primary arrivers, which are capable of killing the larvae and completing their life cycle using the cadaver, and nematodes as secondary arrivers, which cannot kill the larvae themselves but can co-emerge with, or even outcompete, the primary arrivers to exploit the cadavers, probably because of their associated bacteria. In our system, primary arrivers exclude each other due to strong priority effects, while often co-emerging with secondary arrivers. However, our results indicate that priority effects between primary and secondary arrivers can also be strong. Specifically, secondary arrivers may need to reach an insect killed by a primary arriver soon after their arrival to establish themselves in it. Otherwise, if they arrived too late, secondary arrivers would be excluded from the cadaver, as indicated by delayed introduction of secondary arrivers’ symbionts (**Figure 5**).

In conclusion, we suggest that the complex species interactions observed in this study make this insect-parasite system a promising natural microcosm for understanding how priority effects shape communities consisting of both primary and secondary arrivers in host–parasite systems and how their symbionts drive these priority effects (**Figure 6**). The ecological assembly of many other types of biological communities, whether they consist of bacteria, fungi, plants, animals, or any combination of these, are also characterized by priority effects among primary arrivers, among secondary arrivers, and among primary and secondary arrivers. For this reason, we suggest that insect–parasite systems have the potential to inform the basic understanding of how community assembly works more broadly than currently realized.

**Figure 6:**
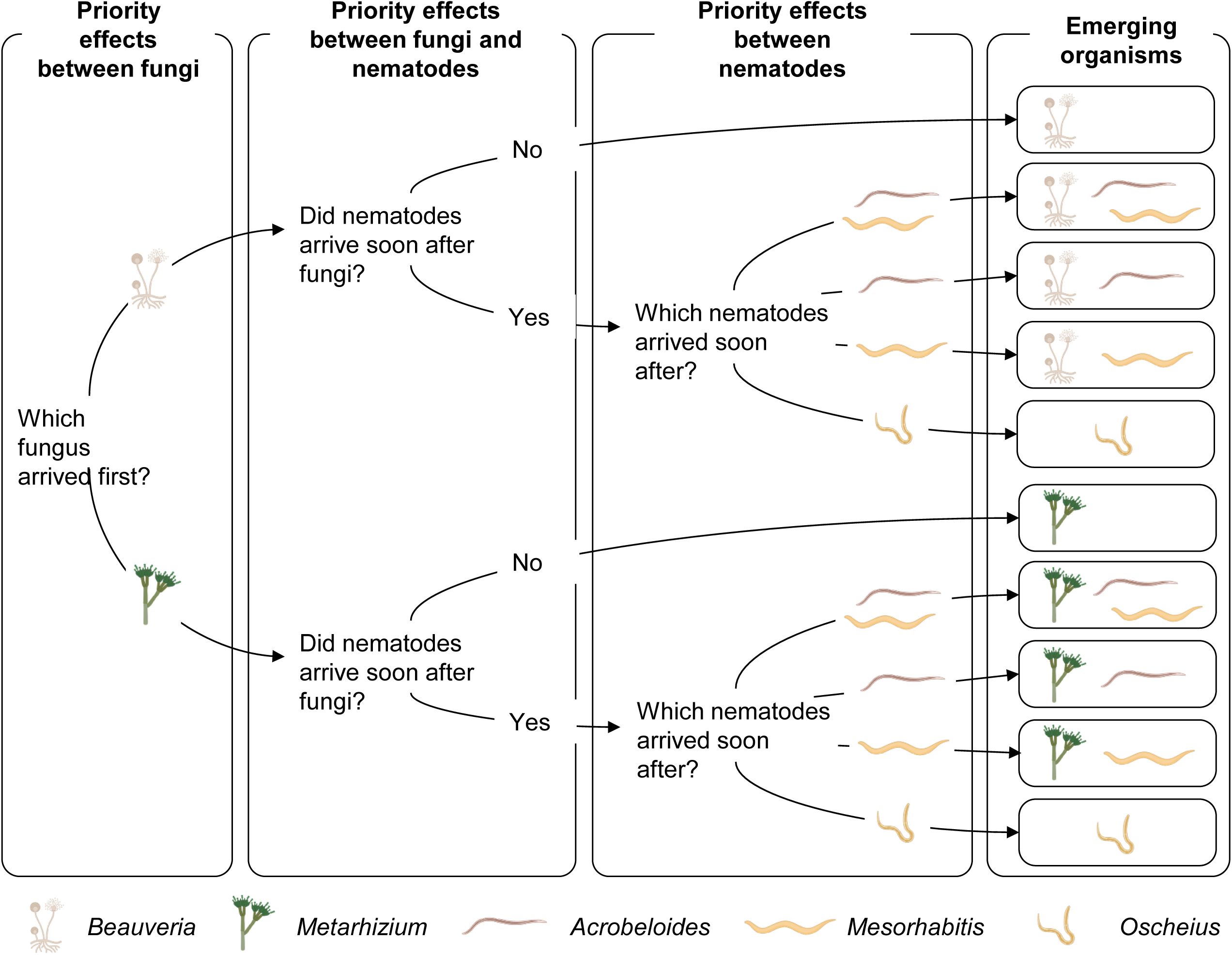
Hypothesis on how priority effects drive fungal and nematode emergence from insect larvae. Findings from this study suggest that fungal and nematode emergence from insect larvae is affected by priority effects among fungi, among nematodes, and among fungi and nematodes.

## Acknowledgments

We thank the members of the Community Ecology Group at Stanford University for comments. Figures were made using R and Biorender.com. This work was funded by the Discovery Grant from the Stanford Doerr School of Sustainability. CJW is an investigator of the Howard Hughes Medical Institute.

---

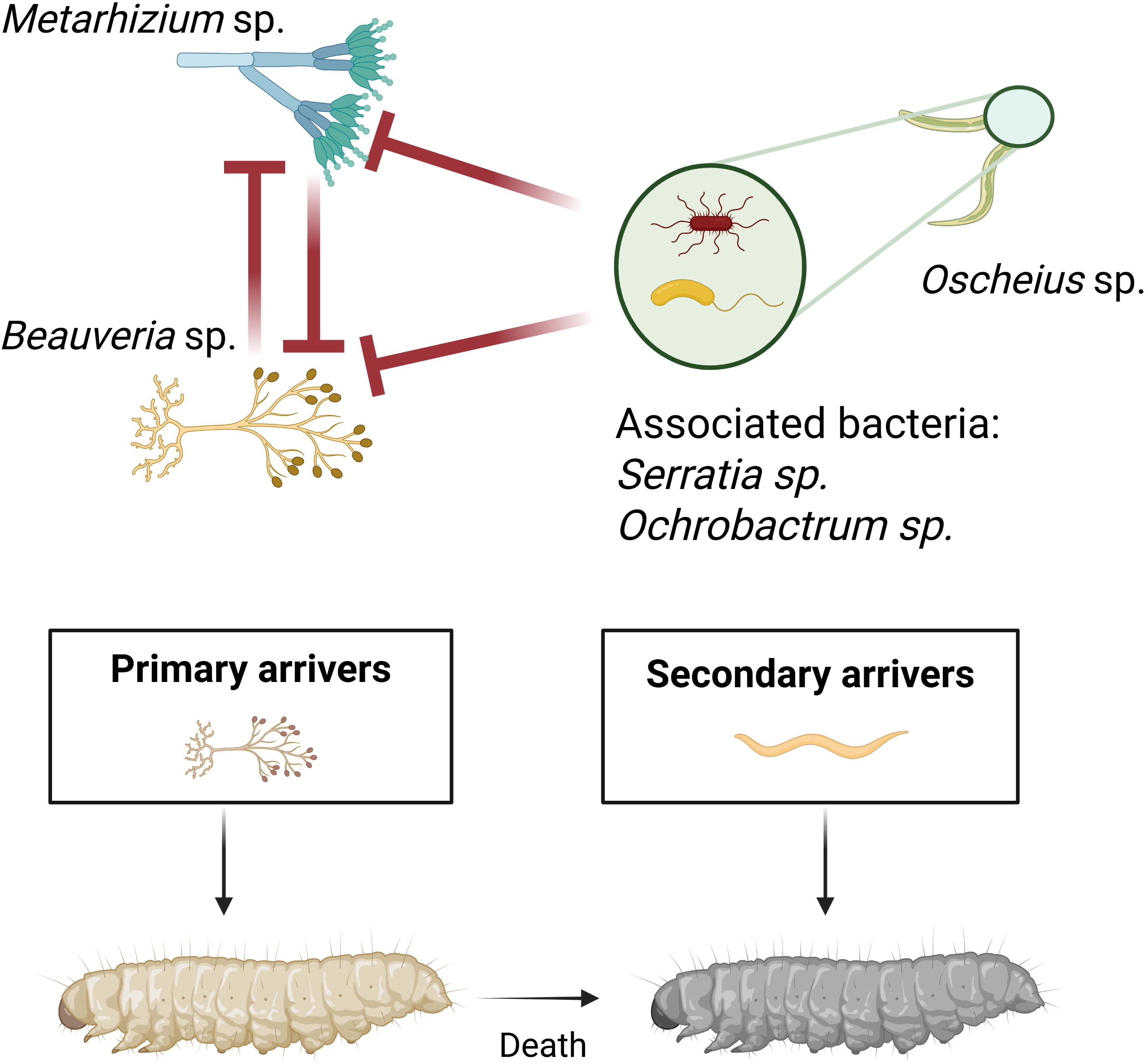

**Figure S1: Fungi that emerged from *Galleria* moth larvae in the field survey.**

Stacked bar plots show monthly survey results for *Metarhizium* and *Beauveria* fungi. Each bar represents the proportion of each fungus for that month. The number of larvae tested for each month is shown above the bars.

**Figure S2: Organisms that emerged from *Galleria* moth larvae placed in the soil collected at location 2.**

Colors and symbols are as in Figure 1, which shows results for location 1.

**Figure S3: Organisms that emerged from *Tenebrio* beetle larvae placed in the soil collected at locations 1 and 2.**

Colors and symbols are as in Figure 1.

**Figure S4: Organisms that emerged from *Tenebrio* beetle larvae in moistened soil from location 2.**

Colors and symbols are as in Figure 1.

**Figure S5: Heatmap of the pairwise Fisher’s exact test on the data shown in Figure 5A**.

Heatmap representing the p value obtained by using the Fisher’s exact test to compare the outcome of *in vitro* co-introduction of *Beauveria* and bacteria. Dark red color show significant difference between the results for the bacterial strain indicated on the x-axis and those for the bacterial strain indicated on the y axis. Asterisks denote statistical significance, where *** means p <0.005, ** means 0.005 < p value <0.01, and * means 0.01 < p <0.05.

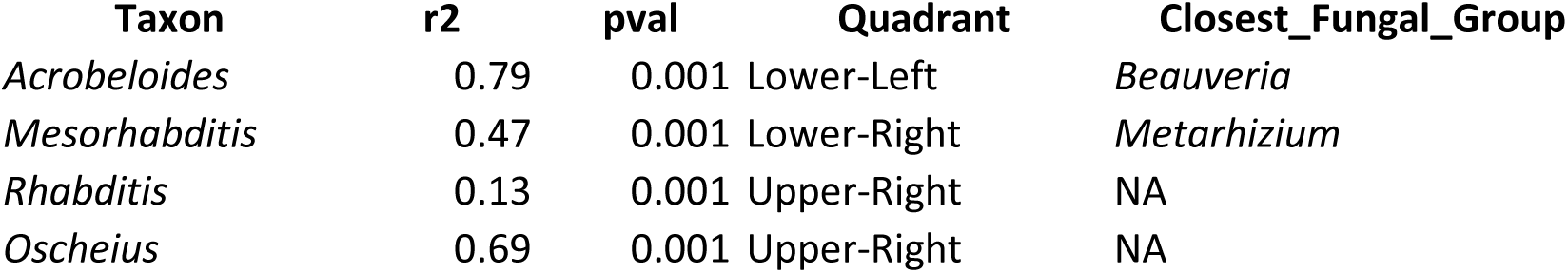

