## Supplementary figures for "Priority effects drive fungal and nematode emergence from insect larvae"

#### Figure S1 : Proportion of *Beauveria* and *Metarhizium* among fungi emerging cadavers

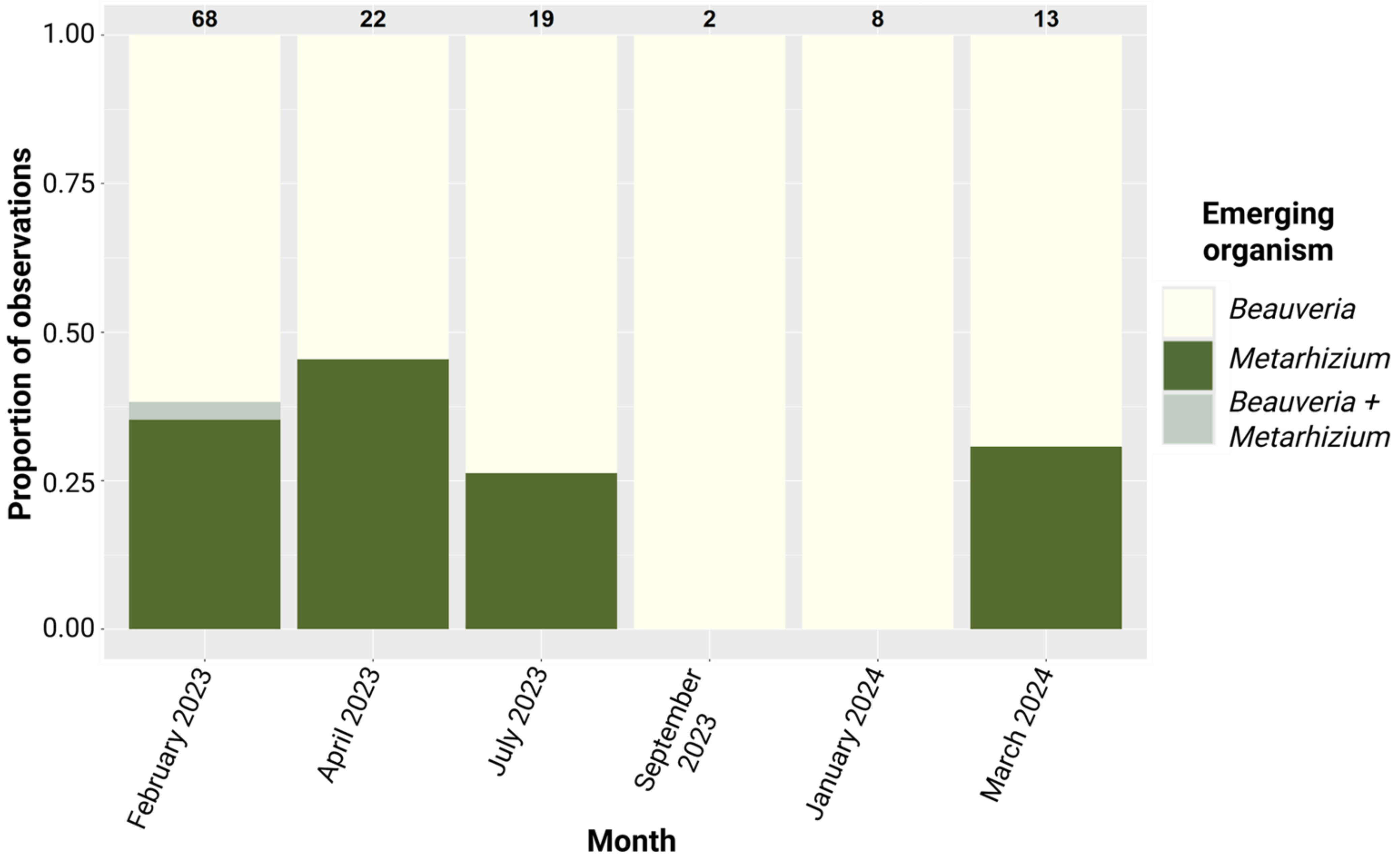

**Figure S2 : Emerging organisms from *Galleria* larva from soil from Location 2**

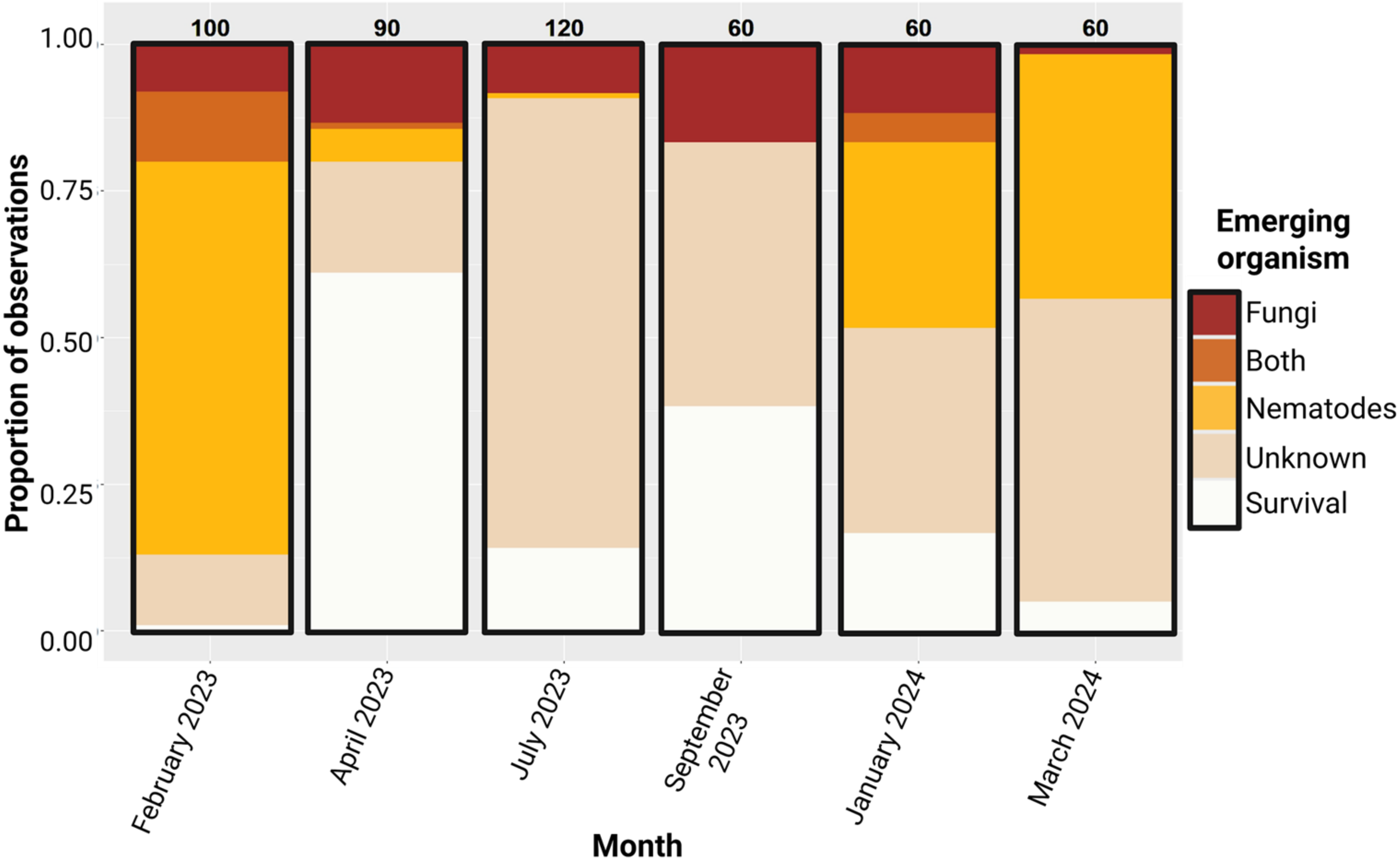

#### Figure S3 : Emerging organisms from *Tenebrio* larva from soil from Location 1 and 2

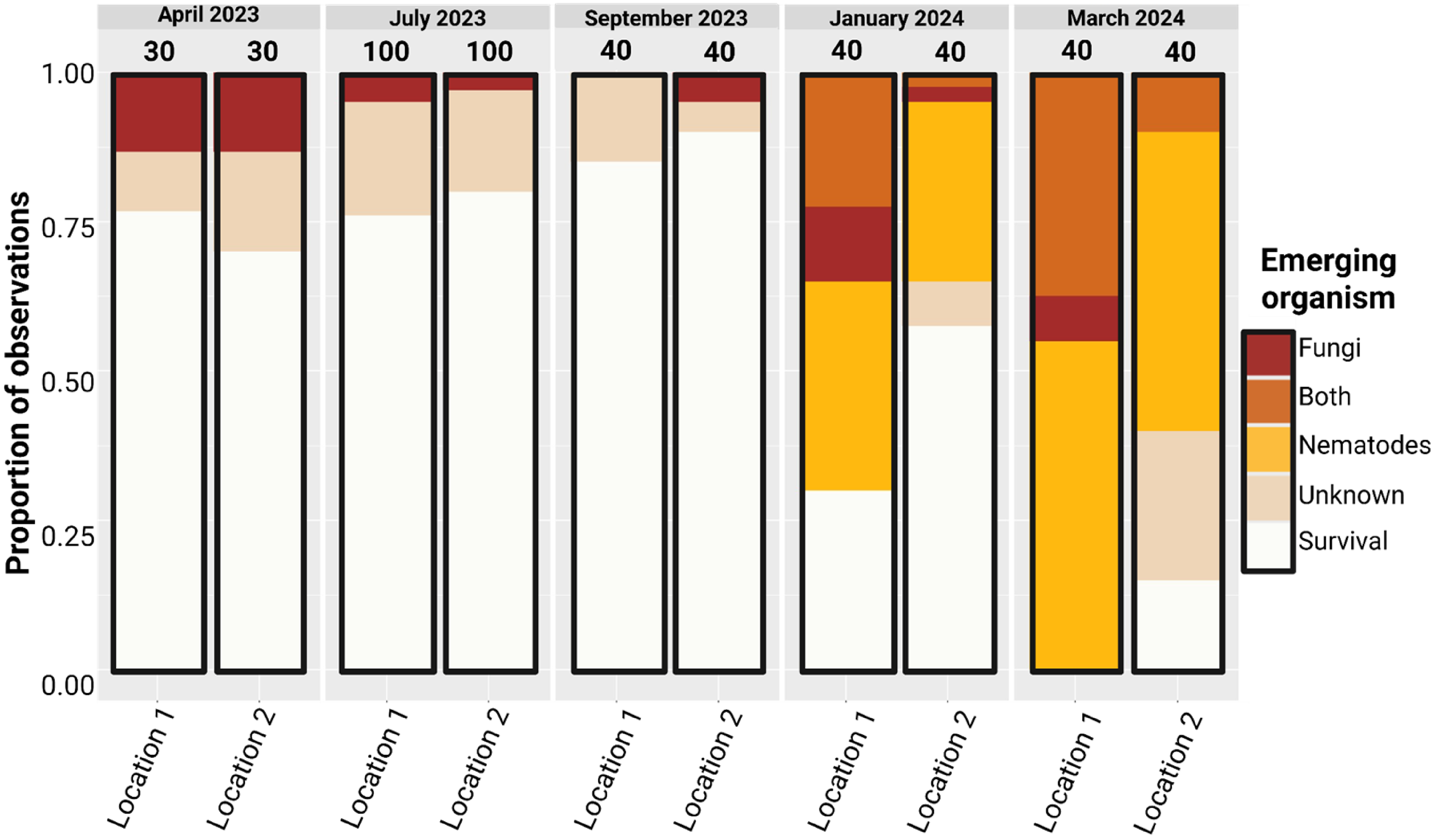

**Figure S4: Emerging organisms from *Tenebrio* larva on moistened soil per month**

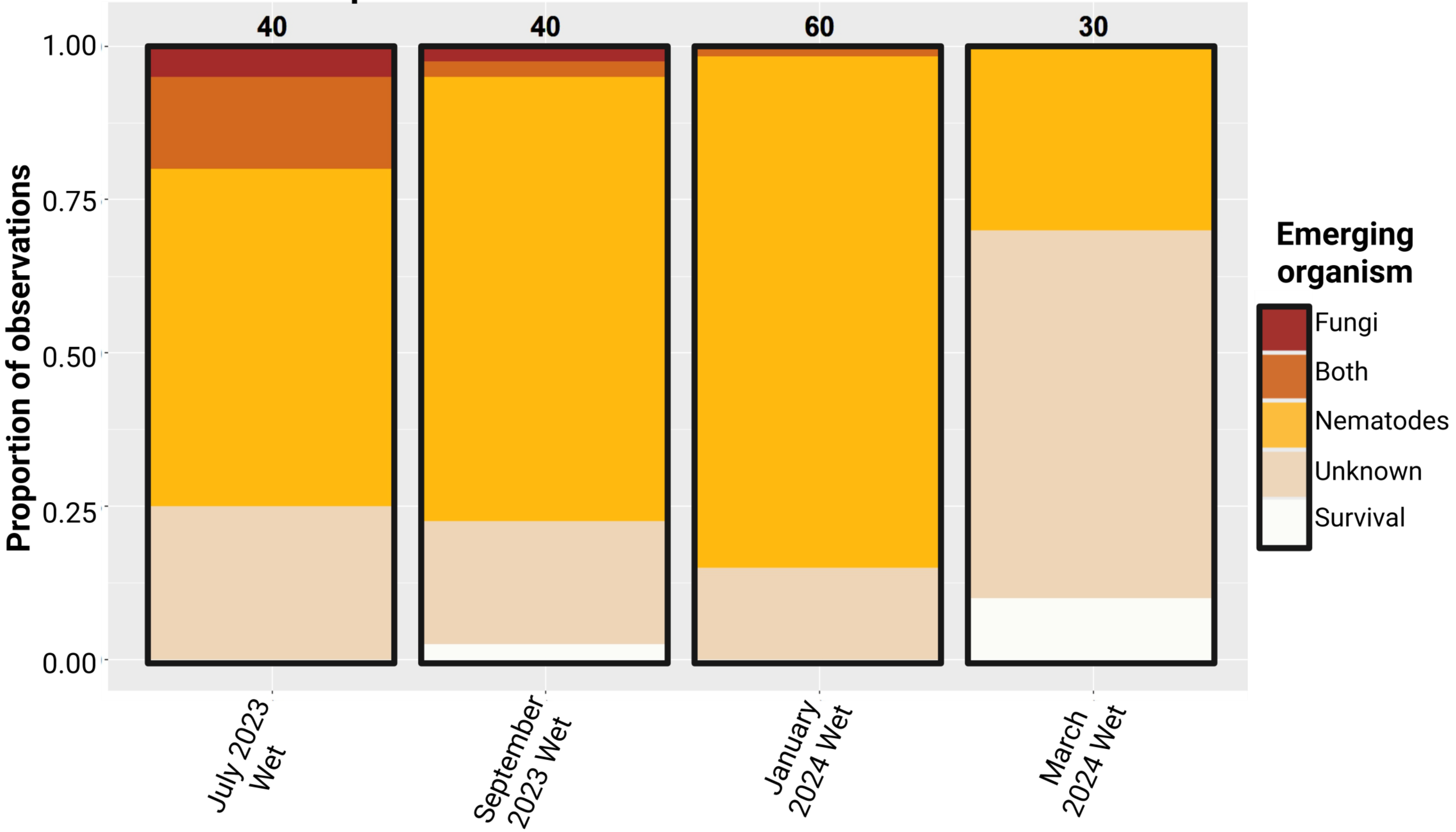

### Figure S5: Heatmap of pairwise Fisher's exact test for Figure 5A

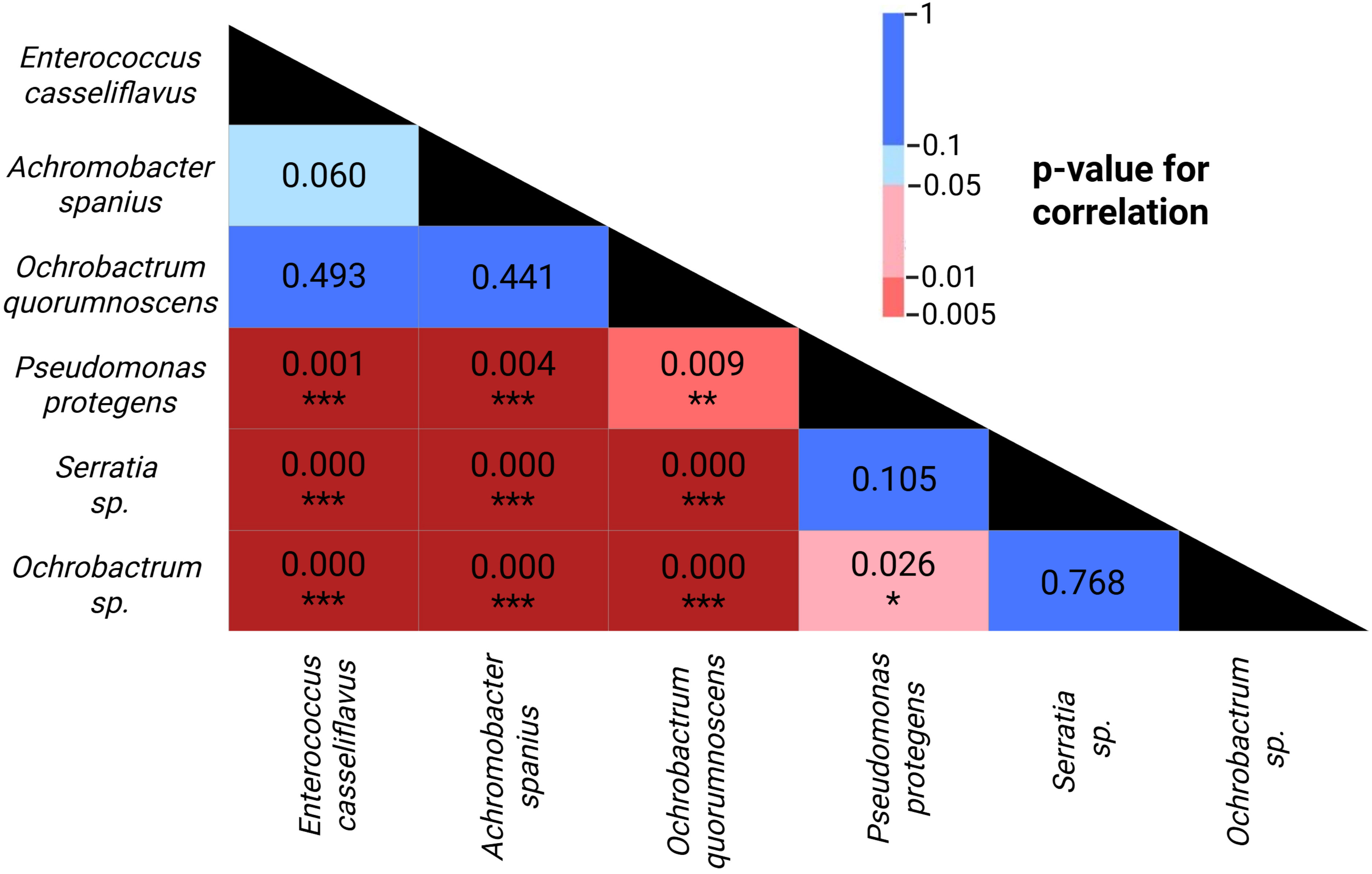
